# Trans-Golgi network-lipid droplet contacts maintain the TGN integrity and function via lipid transfer activities of VPS13B

**DOI:** 10.1101/2020.12.16.423147

**Authors:** Yuanjiao Du, Juan Xiong, Wei-Ke Ji

## Abstract

While the physical interactions between the Golgi apparatus (Golgi) and lipid droplets (LDs) have been suggested through system-level imaging, the bona fide and functional Golgi-LD membrane contact sites (MCSs) remain largely uncharacterized. Here, we demonstrate that vacuolar protein sorting-associated protein 13B (VPS13B) mediates trans-Golgi network (TGN)-LD interactions. VPS13B is specifically accumulated at TGN-LD MCSs with its C-terminal region targeting LDs via an amphipathic helix while a putative WD40 module and a C-terminal Pleckstrin homology (PH) domain independently recognizing TGN via directly binding to Rab6. A putative lipid transfer domain (LTD) at the N-terminal portion of VPS13B binds glycerophospholipids *in vitro*. VPS13B suppression results in severe fragmentation of the TGN, an effect that can be almost completely rescued by the expression of VPS13B-LTD. Collectively, our findings demonstrate that VPS13B mediates lipid transfer at TGN-LD MCSs to maintain TGN integrity and function.

## Introduction

Membrane contact sites (MCSs) are microdomains where opposing membranes of two organelles are tethered by protein–protein or protein–lipid interactions(Eisenberg-Bord et al., 2016; Gatta and Levine, 2017; Lees and Reinisch, 2020; Ugur et al., 2020). A growing body of evidence has supported that MCSs are conserved cellular structures with multiple fundamental functions such as organelle inheritance, fusion and fission, and the transfer of lipids, ions, and other small metabolites(Abrisch et al., 2020; Elbaz and Schuldiner, 2011; Kumar et al., 2018; Lewis et al., 2016; Scorrano et al., 2019; Valverde et al., 2019; Wu et al., 2018). The understanding of MCSs has been greatly improved as tethering, effector and regulatory proteins at MCSs being identified(Eisenberg-Bord et al., 2016; Gatta and Levine, 2017). However, it is clear that only a small portion of the complete cellular puzzle of MCSs have been investigated so far and that therefore new MCSs and key players at such MCSs await discovery(Bohnert and Schuldiner, 2018).

The Golgi apparatus plays a central role in both the anterograde transport of newly synthesized proteins to the plasma membrane and to intracellular organelles, as well as retrograde traffic from endocytic and recycling pathways(Emr et al., 2009; Glick and Nakano, 2009; Malhotra and Mayor, 2006). A typical mammalian Golgi consists of a pile of stapled cisternae (Golgi stacks) that are interconnected by tubules to form the Golgi ribbon. At the entrance site of the Golgi (cis-Golgi), vesicular tubular clusters form the intermediate between endoplasmic reticulum (ER) and the Golgi stack. The exit site of the Golgi (trans-Golgi network, TGN) is the major site for sorting proteins and lipids to distinct cellular locations(Guo et al., 2014). For the efficient modifications, sorting, and trafficking of proteins and membranes, the integrity and structure of TGN must be maintained. However, these mechanisms of TGN maintenance are elusive.

Vacuolar protein sorting-associated protein 13B (VPS13B) is a member of VPS13 family that is highly conserved in animal species. The human genome contains four VPS13 genes (VPS13A/Chorein, VPS13B, VPS13C, and VPS13D)(Velayos-Baeza et al., 2004). A pioneering study has revealed that VPS13A and VPS13C are lipid transporters at ER associated MCSs including endoplasmic reticulum (ER)-mitochondria/LD (VPS13A) and ER-late endosome/lysosome/LD MCSs (VPS13C)(Kumar et al., 2018). VPS13D has been reported to play key roles in the regulation of mitochondrial size and health in *Drosophila*(Anding et al., 2018). Whether VPS13B is associated with a specific type of MCS and what its precise functions at that MCSs may be remains unclear, though VPS13B has been previously implicated in the maintenances of the Golgi structure(Seifert et al., 2011; Seifert et al., 2015) and in cargo transport from early endosomes to recycling endosomes(Koike and Jahn, 2019).

In the present study, we show that VPS13B mediates a novel physical interaction between the TGN and LDs. We further explore the cellular functions of VPS13B mediated TGN-LD MCSs and then investigate the extent to which cellular functions of VPS13B-mediated MCSs are dependent on lipid transfer activities of VPS13B.

## Results and discussion

### VPS13B mediates TGN-LD interactions

System-level spectral imaging has observed that LDs frequently contacted the Golgi apparatus in COS7 cells, suggesting that Golgi apparatus formed MCSs with LDs(Valm et al., 2017). However, the components and cellular functions of such MCSs were unknown. Given that VPS13B associated with Golgi apparatus and might act as a MCSs protein, we hypothesized that VPS13B may function at Golgi-LDs MCSs. To confirm this hypothesis, we initiated with an examination of the cellular localizations of VPS13B within animal cells. We ectopically expressed superfolder green fluorescent protein (sfGFP)-tagged VPS13B (NM152564.6 isoform1) (VPS13B^sfGFP), in which sfGFP was internally tagged to VPS13B at a site shown to preserve yeast VPS13 function(Lang et al., 2015), in HEK293 cells. We used HEK293 cells for this purpose as they are well suited for the transfection and expression of the huge VPS13B^sfGFP construct (∼20 K base pairs in size).

We then investigated the cellular localization of VPS13B^sfGFP in BODIPY 558/568 (magenta, LD marker)-labeled HEK293 cells expressing Halo-Rab6 as a TGN marker by live-cell confocal microscopy. VPS13B^sfGFP localized to Rab6-positive Golgi structures (Fig. 1A), which is in agreement with previous studies (Koike and Jahn, 2019; Seifert et al., 2015). Remarkably, we noted that a substantially higher portion of LDs (∼51%) were closely associated with TGN upon expression of VPS13B^sfGFP (Fig. 1A, B), compared to either pAcGFP1-Golgi (GFP tagged trans-membrane region of B4GALT1 as a general TGN marker) (Fig. S1A) or Halo-Rab6 transfected cells (Fig. S1B), suggesting a potential role of VPS13B in mediating TGN-LD associations. VPS13B^sfGFP was not homogenous on TGN, and appeared to be enriched at contacts between TGN and LDs (Fig. 1A, insets).

**Fig. 1.**
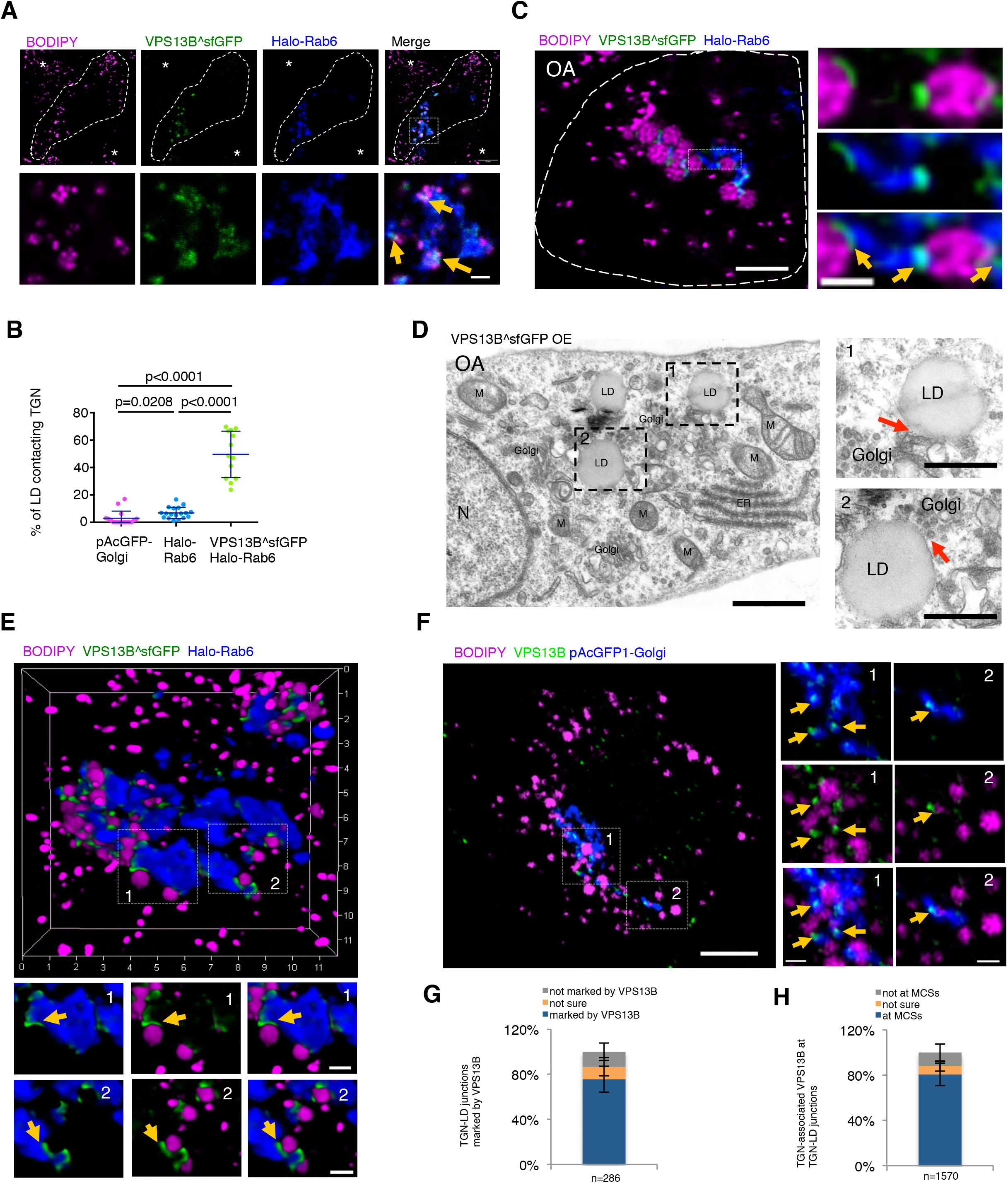
VPS13B localized to TGN-LD MCSs. (A) Representative confocal images of a BODIPY558/568 (LD marker, magenta)-labeled HEK293 cell expressing VPS13B^sfGFP (green), Halo-Rab6 (blue). Yellow arrows denoted the hyper-tethering of LDs to TGN upon VPS13B^sfGFP expression. Asterisks represented neighboring cells with normal distribution of LDs. (B) Quantification of TGN-LD interactions in BODIPY558/568-labeled cells either expressing a general TGN marker pAcGFP-Golgi (n=16), Halo-Rab6 (n=18), or co-expressing VPS13B^sfGFP and Halo-Rab6 (n=12). Mean ± SD. Two tailed unpaired student t test. (C) High-resolution Leica lightning images of a cell as in (A) in medium with OA (500 μM) for 16 hours. Left: whole cell image; right: an inset from a boxed region in whole cell image. Yellow arrows denoted the specific enrichment of VPS13B^sfGFP at TGN-LD MCSs. (D) Representative electron micrographs showing physical interactions between Golgi and LDs (red arrows) in HEK293 cells expressing VPS13B^sfGFP in OA medium with two insets from boxed regions on the right. M: mitochondria; N: nucleus; ER: endoplasmic reticulum. Red arrows denoted potential Golgi-LD MCSs. (E) Top: High-resolution 3D images of a cell as in (A); bottom: two insets from two boxed regions with yellow arrows denoting specific enrichment of VPS13B^sfGFP at TGN-LD MCSs. (F) Airyscan images of fixed BODIPY558/568 (magenta)-labeled HEK293 cells expressing pAcGFP-Golgi (blue) in IF with anti-VPS13B antibody (green). Yellow arrows denoted the specific enrichment of endogenous VPS13B at TGN-LD MCSs. (G) Percentage of TGN-LD junctions (n=286 from 20 cells) marked with endogenous VPS13B. (H) Percentage of TGN-associated, endogenous VPS13B puncta or enrichments (n=1570 from 20 cells) at TGN-LD junctions. Scale bar, 10 µm in whole cell image and 2 µm in insets in (A); 2 µm in whole cell image and 0.5 µm in insets in (C); 1 µm in left panel and 0.5 µm in right panel in (D); 1 µm in (E); and 10 µm in whole cell image and 0.5 µm in insets in (F).

We further validated the localization of VPS13B^sfGFP at TGN-LD MCSs via high-resolution Leica lightning imaging. To better visualize LDs, we cultured the cells in medium containing oleic acid (OA), to increase the size and abundance of LDs. We found that VPS13B^sfGFP was preferentially accumulated at TGN-LD MCSs with LDs being tightly tethered to the TGN (Fig.1C).

Furthermore, we directly examined the interactions between the Golgi and LDs by transmission electron microscopy (TEM). Electron micrographs revealed that the LDs extensively contacted the trans-face of Golgi stacks or Golgi-derived vesicles, likely representing TGN, upon VPS13B^sfGFP expression in HEK293 cells cultured in OA medium (Fig. 1D).

The Golgi structures remained largely peri-nuclear, but occupied a considerable volume in three dimensions (3D). Thereby, we used 3D reconstructions of z-stacks through high-resolution live-cell microscopy to examine the localizations of VPS13B^sfGFP relative to TGN-LD junctions. We observed that VPS13B^sfGFP was specifically enriched at the TGN-LD MCSs in 3D (Fig. 1E). The targeting of VPS13B^sfGFP to the TGN-LD MCSs was accompanied by a substantially increased percentage (∼68%) of LDs tightly associating with TGN, compared to Halo-Rab6-transfected cells, in which a small portion (∼10%) of these LDs were in direct contact with the TGN (Fig.S1C, D), as revealed by co-localization analysis based on maximum intensity projections of z-stacks.

Given the dynamic nature of the TGN and LDs, we tracked the movement of LDs relative to the TGN over time in HEK293 cells expressing VPS13B^sfGFP in OA medium. We found that LDs were tightly associated with the TGN, and these VPS13B-mediated associations between TGN and LDs were very stable over time (Fig.S1E). Collectively, our findings suggest that VPS13B facilitates the tethering between the TGN and LDs.

To avoid the potential artifacts of overexpression, we further explored the localization of endogenous VPS13B by immunofluorescence staining (IF) in HEK293 cells (Fig. 1F). Consistently, endogenous VPS13B mainly localized to the TGN structures with ∼76% of potential TGN-LD junctions were marked by VPS13B (Fig. 1G), while ∼82% of TGN-associated VPS13B puncta (or enrichments) present at TGN-LD junctions (Fig. 1H). However, the fluorescence of endogenous VPS13B at these junctions was almost completely lost upon small-interfering RNAs (siRNAs)-mediated VPS13B suppression (Fig. S2), validating the specificity of the anti-VPS13B antibody in IF. One difference was that only spot-like contacts between the TGN and LDs were observed, confirming that the more extensive TGN-LD MCSs are due to expansions of these contacts as a result of VPS13B overexpression.

We further questioned whether VPS13B is a structural tether by examining the co-localization between the TGN and LDs in controls or VPS13B-depleted cells. The efficiency of siRNAs-mediated VPS13B depletion was further confirmed by immunoblots (Fig. S3A). We found that both the TGN structure and LD density were evidently altered upon VPS13B suppression, and these phenotypes will be described below in detail (Fig. 4A, B). When taking into consideration the number of TGN and LDs, we found that the TGN-LD interactions were significantly reduced (∼64%) upon VPS13B suppression, as revealed by TGN-LD co-localization normalized to the number of TGN and LDs based on maximum intensity projections from 3D reconstructions (Fig. S3B-E). Taken together, our results support that VPS13B is a structural tether that mediates TGN-LD interactions.

**Fig. 2.**
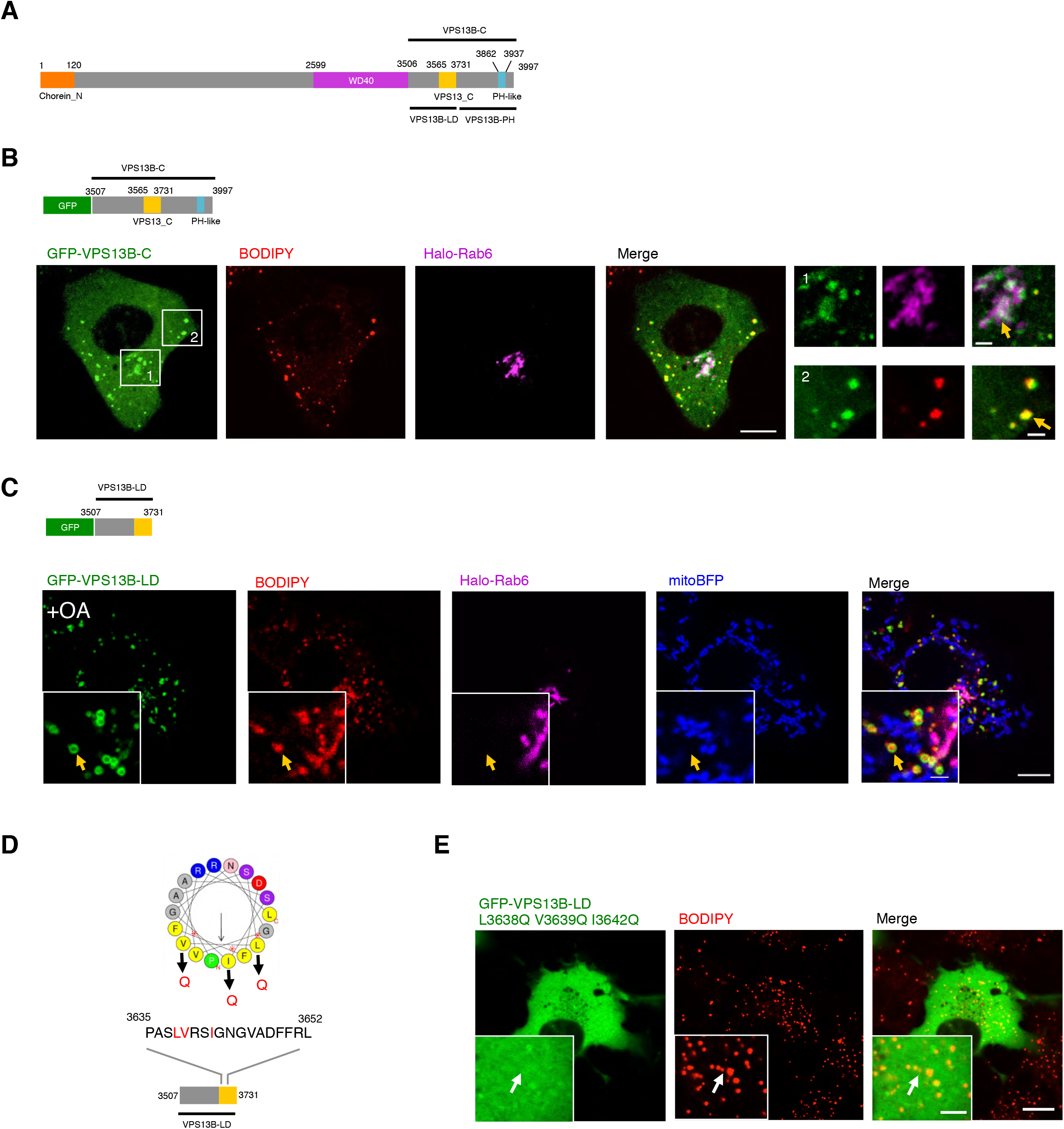
An amphipathic helix in C-terminal of VPS13B target LDs. (A) Diagram of a putative domain organization of VPS13B. (B) Confocal image of a BODIPY558/568 (red)-labeled HEK293 cell expressing truncated mutant GFP-VPS13B-C (green), Halo-Rab6 (magenta), and mitoBFP (blue) with two insets on the right. Yellow arrows indicate GFP-VPS13B-C decorated TGN (top panel) or LDs (bottom panel), respectively. (C) Confocal image of a cell as in (B) expressing truncated mutant GFP-VPS13B-LD (green) and mitoBFP (blue). Yellow arrows indicate that GFP-VPS13B-LD exclusively decorated LDs. (D) Diagram of one predicted amphipathic helical arrangement of residues 3635-3652 (helix-1) of VPS13B via HeliQuest tool(Gautier et al., 2008) (heliquest.ipmc.cnrs.fr). Three point mutations (I/L/V->Q) were introduced to the hydrophobic interfaces of the helix. (E) Confocal image of a BODIPY558/568 (red)-labeled HEK293 cell expressing GFP-VPS13B-LD mutant (green, L3638Q V3639Q I3642Q) with yellow arrows denoting LDs without the decorations of this mutant. Scale bar, 10 μm in whole cell image and 2 μm in insets in (B, C, & E).

**Fig. 3.**
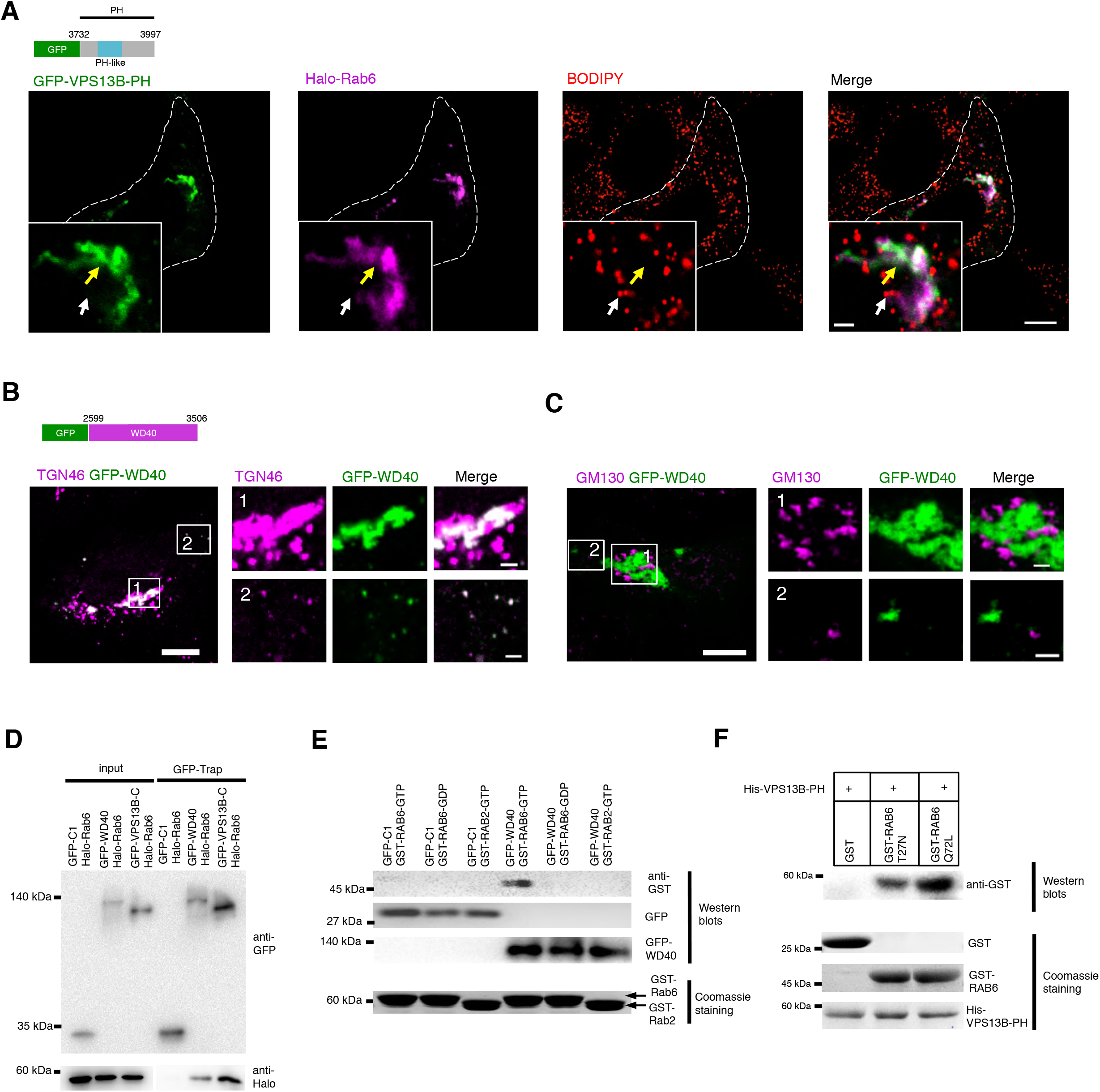
Bipartite interactions between VPS13B and Rab6. (A) Representative confocal images of a BODIPY558/568 (red)-labeled HEK293 cell expressing GFP-VPS13B-PH (green) and Halo-Rab6 (magenta) with an inset on bottom-left. Yellow arrows indicated GFP-VPS13B-PH associating with TGN while white arrows denoting the LDs without GFP-VPS13B-PH. (B) Representative confocal images of a HEK293 cell expressing GFP-WD40 (green) in IF with anti-TGN46 antibody (magenta) with two insets from boxed regions on the right. (C) A HEK293 cell as in (B) was stained with anti-GM130 antibody (magenta) with two insets from boxed regions on the right. (D) GFP-Trap assays of HEK293 cells expressing GFP-VPS13B-C or GFP-WD40 and Halo-Rab6 showing that both GFP-VPS13B-C and GFP-WD40 interacted with Halo-Rab6. (E) Pull-down assays using purified proteins showing that GFP-WD40 bound to GST-Rab6-GTP but not GST-Rab6-GDP or GST-Rab2-GTP *in vitro*. (F) Pull-down assays using purified proteins showing that purified His-VPS13B-PH preferentially bound to purified GST-Rab6-Q72L compared with GST alone, or GST-Rab6-T27N *in vitro*. Scale bar, 10 µm in whole cell image and 2 µm in insets in (A, B).

**Fig. 4.**
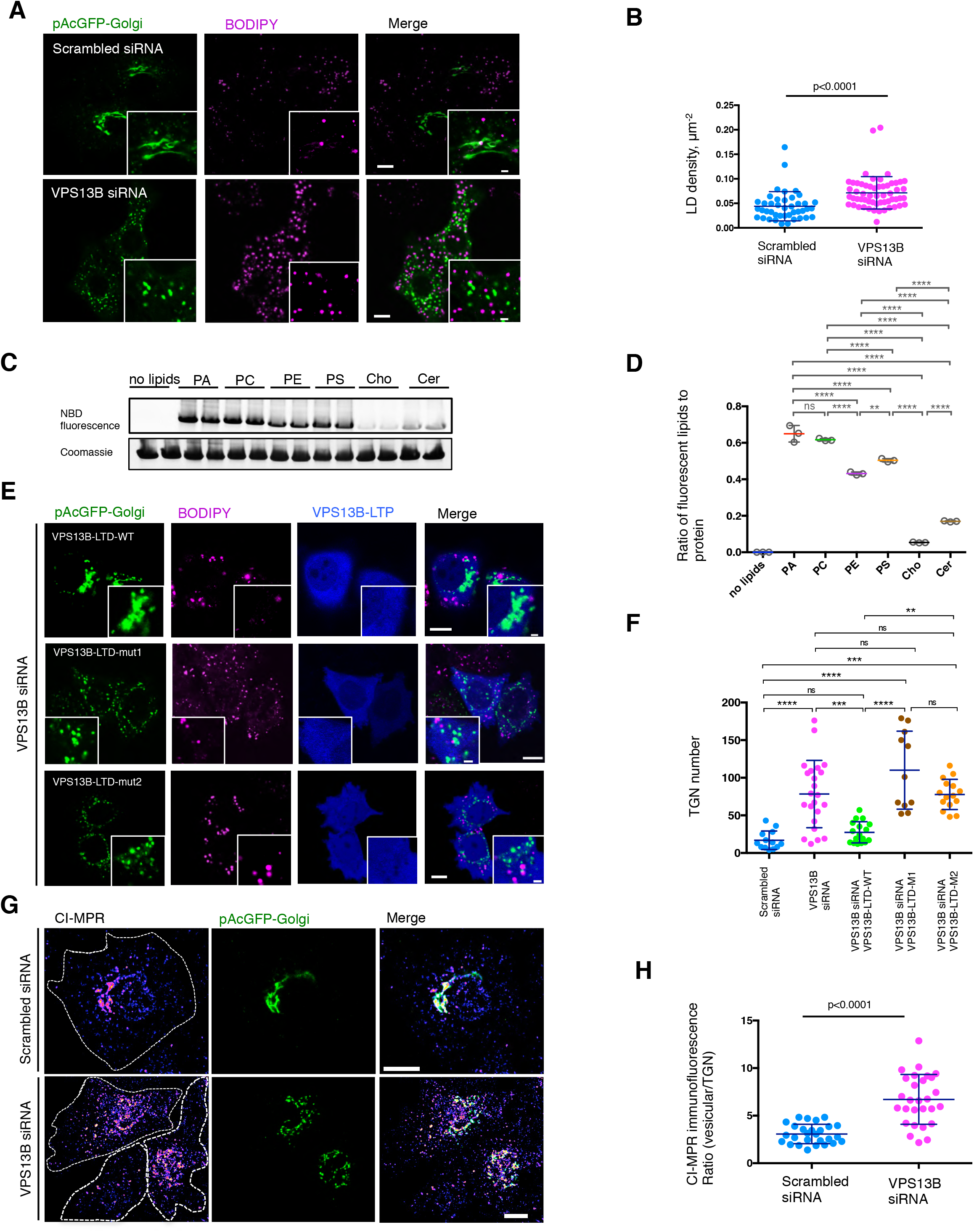
Lipid transfer activity of VPS13B was required for Golgi integrity. (A) Confocal images of BODIPY 558/568 (magenta)-labeled HEK293 cells expressing pAcGFP1-TGN (green) upon treatments with the indicated siRNAs with insets on bottom-right. (B) Quantification of LD density (LD number/cell area) in BODIPY558/568-labeled cells in response to scrambled (n=42) or VPS13B siRNAs (n= 57). Mean ± SD. Two tailed unpaired student t test. (C) *In vitro* lipid-binding assays for VPS13B-LTD. Purified VPS13B-LTD was incubated with NBD-tagged lipids and examined by native PAGE. Phospholipids, visualized by their fluorescence, comigrated with protein, visualized by coomassie blue staining. (D) Ratio of fluorescence of VPS13B-LTD-bound lipids to protein level. P<0.0001 by an ordinary one-way ANOVA followed by Tukey’ s multiple comparisons test. Significant: ****. Mean ± SD (E) Confocal images of BODIPY 558/568 (magenta)-labeled HeLa cells expressing tagBFP2-Golgi (green) and indicated siRNA-resistant VPS13B-LTD constructs (blue) in response to VPS13B siRNAs. (F) Quantification of Golgi number in HeLa cells transfected with scrambled (n=13), VPS13B siRNA (n=23), VPS13B siRNA/VPS13B-LTD-WT (n=17), VPS13B siRNA/VPS13B-LTD-Mut1 (n=11), and VPS13B siRNA/ VPS13B-LTD-Mut2 (n=15). P<0.0001 by an ordinary one-way ANOVA followed by Tukey’s multiple comparisons test. Significant: ****, ***, **. ns: not significant. Mean ± SD (G) Confocal images of internalized anti-CI-MPR antibody by immunostaining in scrambled (top panel) or VPS13B siRNAs (bottom panel) treated HeLa cells expressing pAcGFP-Golgi. (H) The ratio between fluorescence intensity in vesicular structures (outside of the TGN) relative to that of the TGN was measured for scrambled (n=30) or VPS13B siRNAs-treated cells (n=43). Mean ± SD. Two tailed unpaired student t test. Scale bar, 10 μm in whole cell image and 2 μm in insets in (B, F); 10 μm in (H).

### An amphipathic helix in the C-terminal of VPS13B targets LDs

To mediate the tethering of the TGN and LD membranes, VPS13B requires binding sites responsible for recognizing these two membranes. Based on bioinformatics analysis via the Phyre2 structural prediction program, the domain organization of VPS13B consisted of an N-terminal putative lipid transfer domain (LTD) (Chorin_N domain), a putative WD40 module, a VPS13_C domain and a PH-like domain (Fig. 2A)(Bean et al., 2018). We identified that the C-terminal region downstream WD40 module (residues 3507-3997; it was referred to as VPS13B-C hereafter.) was sufficient for recognizing both LDs and Rab6-positive TGN (Fig. 2B). Further molecular dissections demonstrated that a previously uncharacterized region including VPS13_C domain (from residues 3507 to 3731; it was referred to as VPS13B-LD.) mainly targeted LDs with a small portion associating with mitochondria (Fig. S4A), and it was exclusively recruited to LDs upon OA stimulation (Fig. 2C). The region upstream the VPS13_C domain in the VPS13B-LD (residues 3507-3564) was not capable of membrane targeting but instead diffused all over the cytosol (Fig.S4B).

Previous studies have shown that VPS13A and VPS13C targeted LDs through an amphipathic helix(Kumar et al., 2018). To examine whether VPS13B targeted LDs by similar mechanisms, we searched for amphipathic helices in the LD targeting region of VPS13B using the HeliQuest tool(Gautier et al., 2008). We identified an amphipathic helix ranging from residues 3635 to 3652 inside the VPS13_C domain (Fig. 2D). To determine whether this amphipathic helix was required for targeting LDs or not, point mutations were introduced into the hydrophobic interface of the helix (L3638Q V3639Q I3642Q) to break its structure (Fig. 2D). Disruption of this helix blocked the binding of GFP-VPS13B-LD to LDs (Fig. 2E), suggesting that the binding of VPS13B to LDs was mediated through this amphipathic helix.

We noted that the VPS13_C domain alone associated with mitochondria (Fig. S4C), which is in agreement with a prior study (Kumar et al., 2018). Collectively, the C-terminal region of VPS13B (VPS13B-C) contains three membrane-targeting modules: the amphipathic helix (targeting LDs), the VPS13_C domain (targeting mitochondria), and a TGN targeting module (described in detail below). However, our results showed that VPS13B-C targeted LDs and TGN over mitochondria, indicating that the LD/TGN recognition modules outcompeted the VPS13_C domain for membrane targeting. In addition, the presence of multiple membrane-targeting modules in C-terminal of VPS13B suggests that VPS13B localizations may be subjected to regulations depending upon the functional state of the cells as yeast VPS13 localizes at different organelle interfaces depending upon growth conditions(Lang et al., 2015).

### Bipartite interactions between VPS13B and Rab6

Next, we sought to dissect VPS13B to identify the domains that were capable of recognizing the TGN. The C-terminal region containing the putative PH-like domain (residues 3732-3997; it was referred to as VPS13B-PH hereafter) was capable of recognizing the TGN, as revealed by the co-localization between GFP-VPS13B-PH and Halo-Rab6 (Fig. 3A). This result is consistent with a previous study (Seifert et al., 2015). However, PH-like domain alone with GFP tagged at its N-terminal (residues 3862-3937; GFP-PH-like) was diffused in the cytosol without TGN localization upon expression of Halo-Rab6, indicating that the PH-like domain itself is insufficient to target the TGN (Fig. S4D).

Notably, the WD40 module with GFP tagged at its N-terminal (GFP-WD40) co-localized with TGN46 (a TGN marker) (Fig. 3B) but not with the cis-Golgi marker GM130 (Fig. 3C), suggesting that the WD40 module specifically recognized TGN membranes. Additionally, the N-terminal region upstream of the WD40 module (residues 1-2598) was mainly cytosolic (Fig. S4E), which is consistent with its potential role as a long groove to channel lipids between membranes other than membrane targeting modules (Li P, 2020). These results indicated that the WD40 region and VPS13B-PH domain independently recognized the TGN.

We next sought to understand the molecular mechanisms underlying the recognition of the TGN membrane by VPS13B. Since our results, as well as those from previous studies(Koike and Jahn, 2019; Seifert et al., 2015), demonstrated that VPS13B co-localized with Rab6, we evaluated whether either the WD40 region or VPS13B-PH recognized TGN through interacting with Rab6. Indeed, interactions between the WD40 module and Rab6 were confirmed by GFP-Trap assays in HEK293 cells (Fig. 3D). In addition, GFP-Trap assays further verified interactions between a C-terminal region containing VPS13B-PH (VPS13B-C) and Rab6 (Fig. 3D), which is in agreement with a previous study (Seifert et al., 2015). Our results suggest that the WD40 module and VPS13B-PH independently recognizes the TGN through Rab6.

To confirm this hypothesis, we investigated whether these two TGN-targeting modules directly bind to Rab6 *in vitro*. At this time we were unable to produce purified WD40 in sufficient quantities for *in vitro* pull-down assays. Alternatively, we used GFP-Trap assays to pellet GFP-WD40 from HEK293 cells expressing GFP-WD40 using high-salt (500 mM NaCl) lysis buffer. After washing out proteins that could co-pellet with GFP-WD40 under high-salt conditions, GFP-WD40 beads were incubated with purified Glutathione S-transferase (GST)-Rab6-GTP, GST-Rab6-GDP, or GST-Rab2-GTP, respectively, and then examined via western blots. Indeed, GFP-WD40, but not the GFP tag, bound to Rab6-GTP but neither Rab6-GDP nor Rab2-GTP (Fig. 3E). This result indicates that GFP-WD40 directly binds the active Rab6. In addition, *in vitro* pull-down assays with purified proteins showed that 6xHis tagged VPS13B-PH interacted directly with constitutive-active mutant Rab6 Q72L, and exhibited less interaction with dominant-negative mutant Rab6 T27N, but showed no interaction with GST tag (Fig. 3F). Taken together, our results demonstrate that the WD40 module and VPS13B-PH independently bind the active form of Rab6, and indicate that VPS13B is a RAB6 effector via these domains. The bimodal interactions between VPS13B and Rab6 are consistent with a loop structure at the C-terminal of VPS13 proteins, in which the WD40 region and the VPS13B-PH domain might be in proximal to each other (Fig. S6A) (De et al., 2017) (Li P, 2020).

### Lipid transfer activities of VPS13B are required for TGN integrity and function

We next investigated the cellular functions of VPS13B-mediated TGN-LD MCSs by examining the morphologies of the TGN and LDs upon siRNAs-mediated VPS13B depletion. We used HeLa cells for this purpose, as the expression level of VPS13B in HeLa is substantially higher than that in HEK293 cells (data not shown). We found that the TGN was evidently fragmented in VPS13B-depleted HeLa cells (Fig. 4A). In addition, VPS13B silencing led to a substantial increase (∼63%) on LD density (LD number/cell area) (Fig. 4B). This finding indicates essential roles of VPS13B in the maintenance of the TGN structure and the cellular LD density, the former of which is in accord with previous studies. (Seifert et al., 2011).

The localization of VPS13B at TGN-LD MCSs was consistent with its potential role in lipid transfer between TGN and LDs. We therefore investigated the extent to which lipid transfer activities of VPS13B were responsible for TGN integrity. We initiated by directly testing the possibility that VPS13B might bind lipids. We were unable to produce sufficient quantities of full-length VPS13B proteins for lipid binding analysis. Thereby, we used purified LTD (residues 1-345) of VPS13B for this purpose.

We examined the lipid-binding activities of VPS13B-LTD via *in vitro* lipids binding assays. VPS13B-LTD co-migrated with nitrobenzoxadiazole (NBD)-labeled glycerophospholipids: phosphatidylserine (PS), phosphatidylcholine (PC), phosphatidylethanolamine (PE) and phosphatidic acid (PA), to a substantial lesser extent with cholesterols or ceramides, as assessed by native gel (Fig. 4C, D), a phenotype similar to yeast VPS13(Kumar et al., 2018; Li P, 2020) or ATG2(Valverde et al., 2019), both of which were highly conserved with VPS13B in LTD. As a control, we constructed two VPS13B-LTD mutants, in which multiple conserved hydrophobic residues were mutated to hydrophilic residues based on sequence alignment among N-terminal regions of VPS13B, ATG2A/B, and yeast VPS13 (Fig. S5A). These mutated residues acted as a band to block lipid transfer through the groove (Li P, 2020; Valverde et al., 2019).

Remarkably, the TGN fragmentation induced by VPS13B depletion was almost completely rescued by siRNA-resistant wild-type (WT) VPS13B-LTD (Fig. 4E, F). In contrast, neither of the VPS13B-LTD mutants was able to rescue the TGN defects (Fig. 4D, E), despite their expression levels being comparable to the WT VPS13B-LTD (Fig. S5B). This finding indicates that the lipid transfer activity of VPS13B is fully sufficient and required for the maintenance of TGN structures.

We further validated whether the TGN fragmentation induced by VPS13B suppression impaired cargo sorting via tracking the cellular transport of CI-MPR (cation-independent mannose 6-phosphate receptor). In control cells, the CI-MPR antibody was internalized with the receptor and accumulated at the TGN (Fig. 4G, top panel). In contrast, cells that were depleted for VPS13B showed fragmented TGN, along with a significant reduction (∼2-fold) in CI-MPR fluorescence at the TGN, concomitant with increased levels of fluorescence at non-TGN peripheral vesicles (Fig. 4G, bottom panel; Fig. 4H). The overall levels of CI-MPR in control cells versus VPS13B-silenced cells remained consistent (Fig. S5C). This finding suggests that VPS13B-mediated TGN integrity is required for efficient CI-MPR cargo sorting.

Taken together, these results revealed critical roles of VPS13B in both the TGN integrity/function and cellular LD density, consistent with its role as a lipid transporter at TGN-LD MCSs. Our findings further suggest that VPS13B may mediate bulky lipid transfer from LDs to the TGN to support TGN integrity and function (Fig. S6B). The mechanisms of VPS13B-mediated directional lipid transport warrant further investigations. In considering the contribution of lipid transfer to TGN integrity and function, we envision a potential role of LDs as a reservoir of lipid fatty acids moieties to support the lipid homeostasis of TGN membranes.

The discovery of the novel TGN-LD MCSs and the essential protein VPS13B that transfers lipids at these MCSs to ensure TGN integrity and function broadens the scope of our understanding of MCSs. Loss-of-function mutations of VPS13B leads to Cohen disease, a complex syndrome featured by global developmental delay and intellectual disability (Kolehmainen et al., 2004), and our discoveries will provide mechanistic insights on this disease.

## METHODS AND MATERIALS

### Plasmids and siRNA oligonucleotides

Human VPS13B ORF was cloned from HeLa cDNA, and fully sequenced (NM152564.6. isoform1). The internal sfGFP tagged VPS13B (VPS13B^sfGFP) was generated by inserting sfGFP sequence after V1265 of VPS13B in frame. All of truncated VPS13B mutants used in this study were generated by PCR using VPS13B^sfGFP as template and cloned it into mGFP-N1 (addgene 54767) or mGFP-C1 (addgene 54579). The VPS13B-LTD mutants were synthesized by Genscript Biotech, and then cloned to GFP-C1 vector. pET28a-6xHis-SUMO-VPS13B-LTD was generated by inserting VPS13B residues 1–345 coding sequence obtained by PCR using VPS13B^sfGFP as template into pET28a-6xHis-SUMO vector, which was generated by inserting 6xHis and sumo tag into pET28a vector. pAcGFP-Golgi was bought from Clontech (Cat. No.632464). Halo-Rab6 was generated by inserting Rab6A coding sequence obtained by PCR using HeLa cDNA as template into Halo-C1 vector. mito-BFP was a gift from Gia Voeltz (Addgene 49151), and was previously described(Friedman et al., 2011). All of constructs used in this study were generated by using ClonExpress®II One Step Cloning kit (Vazyme, C112-01).

Oligonucleotides for human VPS13B siRNAs were synthesized by RiboBIO (Guangzhou, CN) against target sequence 5’-GACCTTACTTGTCATAATA-3’ (siRNA#1); 5’-GCCTATGTTTATTCGTATA-3’ (siRNA#2); As a control, scrambled siRNA 5’-CGUUAAUCGCGUAUAAUACGCGUAT-3′ (RiboBIO) was used.

### Antibodies and reagents

Anti-GFP (AE011, Abclonal), anti-Halo (G9211; Promega), anti-Tubulin (100109-MM05T; Sinobiological) were used at 1:1000 dilutions for Western blot. Anti-VPS13B (HPA043865; Sigma), anti-TGN46 (13573-1-AP; Proteintech), anti-GM130 (11308-1-AP; Proteintech), anti-CI-MPR (A3762; Abclonal) antibodies were used 1:100 for immunofluorescence.

The following reagents were used in this study: Oleic acid (O1008; sigma); BODIPY 558/568 (ThermoFisher, D3835), and antibiotics such as G418 (10131027) and puromycin (A1113803) were obtained from Thermofisher. All EM reagents were purchased from Electron Microscopy Sciences. Lipids were purchased from Avanti Polar Lipids: NBD-PC (810133), NBD-PE (810144), NBD-PS (810198), NBD-PA (810138), NBD-ceramide (810211), and NBD-Cholesterol (810250).

### Cell culture and transfection

Human embryonic kidney 293 cells (ThermoFisher, R70507), human cervical cancer HeLa cell line (ATCC), and Human embryonic kidney 293T cells (ATCC) were grown in DMEM (Invitrogen) supplemented with 10% fetal bovine serum (Gibco) and 1% penicillin/streptomycin. All of the cell lines used in this study are free of mycoplasma contamination.

For transfection, cells were seeded at 4 × 10^5^ cells per well in a six-well dish 16 h before transfection. Plasmid transfections were performed in OPTI-MEM (Invitrogen) with 2 µl Lipofectamine 2000 per well for 6 h, followed by trypsinization and replating onto glass-bottom confocal dishes at 3.5 × 10^5^ cells per well. Cells were imaged in live-cell medium (DMEM with 10%FBS and 20 mM Hepes no antibiotics) 16–24 h after transfection. For all transfection experiments in this study, the following amounts of DNA were used per 3.5cm well (individually or combined for cotransfection): 1000 ng for VPS13B^sfGFP; 500 ng for truncated VPS13B mutations; 500 ng for pAcGFP-Golgi; 50 ng for Halo-Rab6 or Rab6 mutants; 500 ng for mito-BFP. For siRNA transfections, cells were plated on 3.5cm dishes with 30–40% density, and 2 µl lipofectamine RNAimax (Invitrogen) and 50 ng siRNA were used per well. 48 hours after transfection, second round of transfection was performed with 50 ng siRNAs. Cells were analyzed 24 h post-second transfection for suppression.

### BODIPY 558/568 staining in live cell

Cells were washed once with PBS, and was changed to complete medium containing 3 µM BODIPY558/568, and incubated at 37°C for indicated period of time. Cells were washed with PBS three times and changed to imaging medium (DMEM supplemented with 10% FBS and 20 mM Hepes without phenol red) prior to imaging.

### Halo staining in live cell

Cells were incubated with complete medium with 5nM Janilia Fluo® 646 HaloTag® Ligand for 30 minutes. Cells were washed three times with complete medium to remove extra ligands, followed by incubation for another 30 minutes. Medium was replaced with imaging medium to remove unconjugated Halo ligands that has diffused out of the cells prior to imaging.

### Live imaging by confocal microscopy

Cells were grown on glass-bottom confocal dishes. Confocal dishes were loaded to a laser scanning confocal microscope (LSM780, Zeiss, Germany) equipped with multiple excitation lasers (405nm, 458nm, 488nm, 514nm, 561nm, 633nm) and spectral fluorescence GaAsP array detector. Cells were imaged with the 63× 1.4-NA iPlan-Apochromat 63x oil objective using the 405-nm laser for BFP, 488-nm for GFP, 561-nm for mStrawberry, OFP, tagRFP or mCherry. Cells were imaged in live cell chamber supplied with 5% CO^2^ at 37°C.

### High-resolution airyscan live cell imaging

Cells on confocal dishes were loaded to a laser scanning confocal microscope (LSM900, Zeiss, Germany) with Airyscan2 in equipped with ≥ 32 GaAsP-PMT array detector, 63x/1.4 oil objective (working distance 190 µm), four lasers (405nm 5mW, 488nm 5mW, 561nm 5mW, 640nm 5mW) with corresponding filters. Cells were imaged with 405-nm laser for BFP, 488-nm for GFP, 561-nm for OFP, tagRFP or mCherry and 640-nm for Janilia Fluo® 646 HaloTag® Ligand.

### Live cell imaging by Leica SP8 equipped with lightning super-resolution module

Cells on confocal dishes were loaded to Leica SP8 equipped with lightning super-resolution module equipped with HC PL APO CS2 100x/1.4 oil objective, four lasers (415nm, 499nm, 567nm, and 662nm) with corresponding filters. Cells were imaged in optiMEM with Hepes buffer in live cell chamber supplied with 5% CO_2_ at 37°C.

### Immunofluorescence staining

Cells were fixed with 4% PFA (paraformaldehyde, Sigma) in PBS for 10 min at room temperature. After washing with PBS three times, cells were permeabilized with 0.1% Triton X-100 in PBS for 15 min on ice. Cells were then washed three times with PBS, blocked with 0.5% BSA in PBS for 1 h, incubated with primary antibodies in diluted blocking buffer overnight, and washed with PBS three times. Secondary antibodies were applied for 1 h at room temperature. After washing with PBS three times, samples were mounted on Vectashield (H-1000; Vector Laboratories).

### Electron microscopy

OA-stimulated HEK293 cells were fixed with 2.5% glutaraldehyde in 0.1M Phosphate buffer, pH7.4 for 2h at room temperature. After washing three times with 0.1M Phosphate buffer, cells were scraped and collected with 0.1M phosphate buffer followed by centrifugation at 3000 rpm. The pellet was resuspended in PBS (0.1M), and centrifuged at 3000rpm for 10min. This step was repeated three times. The samples were post-fixed with pre-cold 1% OsO4 in 0.1M Phosphate buffer for 2-3 hour at 4°C, followed by rinsing with PBS 3 times (3×20 min). The samples were dehydrated in graded ethanol (50%, 70%, 85%, 90%, 95%, 2×100%) with 15 min for each condition. The penetrations were performed in an order of acetone-epoxy (2:1); acetone-epoxy (1:1); epoxy. Each round of penetration was performed at 37°C for 12 hours. The samples were embedded in epoxy resin using standard protocols. Sections parallel to the cellular monolayer were obtained using a Leica EM UC7 with the thickness of 60-100 nm and examined under HT7800/HT7700. Golgi apparatus and LDs were identified based on their respective morphology and were traced by hand. Golgi apparatus contacting LDs within 30 nm at any point on the Golgi circumference was identified as potential contact sites.

### GFP-trap assay

GFP-Trap (GFP-trap agorose beads, ChromoTek) was used for detection of protein-protein interactions and the GFP-Trap assays were performed according to the manufacturer’s protocol. 5% input was used in GFP traps unless otherwise indicated.

### *In vitro* Pull-down assays of GFP-WD40 and GST-Rab6

HEK293 cells transiently transfected with GFP-only or GFP-WD40 were lysed in high-salt lysis buffer (RIPA buffer containing 500 mM NaCl, proteasome inhibitors and PMSF). GFP-Trap was used to pellet GFP-only or GFP-WD40 from cell lysate, followed by washing with high-salt lysis buffer for 10 times.

Purified GST-Rab6 or GST-Rab2 was incubated with 100 µM GTP or GDP at 25°C for 30min, followed by addition of 30 mM MgCl2 and incubation at 25°C for another 1 h. Finally, GTP or GDP-loaded GST-Rab6 or GST-Rab2 was subjected to GFP pull-down assays using GFP-WD40 or GFP-only loaded GFP-Trap beads overnight at 4°C, respectively. The GFP-Trap beads were washed with freshly prepared HNM buffer (20 mM Hepes, pH 7.4, 0.1 M NaCl, 5 mM MgCl2, 1 mM DTT, 0.2% NP-40). The GFP-Trap beads were re-suspended in 100 µl 2x SDS-sampling buffer. Re-suspended beads were boiled for 10 min at 95°C to dissociate protein complexes from beads. Western blots were performed using anti-GST antibodies and anti-GFP antibodies. The coomassie staining was performed for purified GST-Rab6 and GST-Rab2 as loading controls.

### *In vitro* Pull-down assays of His-VPS13B-PH and GST-Rab6 mutants

Purified GST-only, GST-Rab6 Q72L or GST-Rab6 T27N was incubated with 20 µl Glutathione resin (L00206 GenScript) at 4°C overnight, respectively, followed by washing beads with freshly prepared HNM buffer (20 mM Hepes, pH 7.4, 0.1 M NaCl, 5 mM MgCl_2_, 1 mM DTT, 0.2% NP-40). Purified His-VPS13B-PH was subjected to GST pull-down using GST-only, GST-Rab6 Q72L or GST-Rab6 T27N loaded glutathione resin overnight at 4°C, respectively. Glutathione resin was washed with freshly prepared HNM buffer (20 mM Hepes, pH 7.4, 0.1 M NaCl, 5 mM MgCl2, 1 mM DTT, 0.2% NP-40). Glutathione resin was re-suspended in 100-µl 2x SDS-sampling buffer. Re-suspended resin was boiled for 10 min at 95°C to dissociate protein complexes from beads. Western blots were performed using anti-His antibodies. The coomassie staining was performed for purified GST-only, GST-Rab6 Q72L, GST-T27N and purified His-VPS13B-PH as loading controls.

### Protein expression and purification

Proteins of interest were expressed in TSsetta (DE3) Chemically Competent Cell. Cells were grown at 37°C to an OD_600_ of 0.6–0.8. Protein expression was induced with 0.1 mM IPTG, and then cells were cultured at 16°C for another 16 h. Cells were pelleted, resuspended in buffer A (20 mM Tris-Hcl, pH 8.0, 300 mM NaCl, 10 mM imidazole, 1mM DTT and 10% glycerol) supplemented with protease inhibitors (TargetMoI) and lysed in a JY88-IIN cell disruptor. Cell lysates were centrifuged at 14,000 g for 30 min. Supernatant was incubated with Ni-NTA resin (QIAGEN) for 2 h at 4°C, and then the resin were passed through via gravity flow. Resin was washed with 5 bed volumes of buffer B (20 mM Tris-Hcl, pH 8.0, 300 mM NaCl, 20 mM imidazole, 1× protease inhibitor cocktail (TargetMoI)), followed by 5 bed volumes of buffer C (20 mM Tris-Hcl, pH 8.0, 300 mM NaCl, 50 mM imidazole, 1× protease inhibitor cocktail (TargetMoI)) and 5 bed volumes of buffer D (20 mM Tris-Hcl, pH 8.0, 300 mM NaCl, 100 mM imidazole, 1× protease inhibitor cocktail (TargetMoI)). Retained protein was eluted from the resin with buffer B supplemented with 200 mM imidazole and 500 mM imidazole respectively. The eluted protein solutions were concentrated in a 3-kDa molecular weight cutoff (MWCO) Amicon centrifugal filtration device.

### *In vitro* lipid-binding assay

19 µl purified VPS13B-LTD (1.5 mg/ml) was mixed with 1 µl of either NBD-labeled PA, PC, PE, PS, ceramide, or Cholesterol in 20-µl total reaction volumes and incubated at 4°C for 3 h. Samples were loaded onto 12% Precast Native gels with Hepes-Tris buffer and run for 4 h at 90 V on ice. NBD fluorescence was visualized using a Bio-rad ChemiDoc XRS+ 170-8265. Then gels were stained with Coomassie blue G250 to visualize total protein.

### CI-MPR uptake and trafficking analysis

CI-MPR uptake and trafficking assays were performed as described previously (Hoyer et al., 2018). Briefly, HeLa cells transiently transfected with pAcGFP-Golgi were incubated with 20 µg/µL rabbit anti-CI-MPR monoclonal antibody (Abclonal A3762) in serum-free DMEM for 1 hour, rinsed with PBS, fixed with 4% Paraformaldhyde, solubilized with Triton-X and immunostained with fluorescent anti-rabbit antibodies 1:500. The fluorescence intensity of whole cell area and a region encompassing TGN was measured by FIJI. Vesicular fluorescence was obtained by subtracting the TGN region fluorescence intensity from whole cell fluorescence. A ratio of vesicular fluorescence/Golgi fluorescence was calculated for each cell.

### Image analysis

All image analysis and processing was performed using ImageJ (National Institutes of Health). Colocalization-based analysis of TGN-LD MCSs was performed by colocalization plugin (imageJ, NIH) with the following settings: Ratio (0-100%): 50; Threshold channel 1(0-255): 50; Threshold channel 2 (0-255): 50; Display value (0-255): 255. MCSs were automatically identified by colocalization plugin with overlapping pixels representing potential MCSs. LD or Golgi number was measured manually with assistance of an imageJ plugin Cell Counter.

### Statistical analysis

All statistical analyses and p-value determinations were performed in GraphPad Prism6. All the error bars represent mean ± SD. To determine p-values, ordinary one-way ANOVA with Tukey’s multiple comparisons test were performed among multiple groups and a two-tailed unpaired student t-test was performed between two groups.

## Acknowledgements

We thank Marianna Leonzino (Yale University School of Medicine) and Karin M. Reinisch (Yale University School of Medicine) for valuable advices with transfections and expressions of VPS13 constructs in mammalian cells. We thank Guangjun Cai (Huazhong University of Science and Technology) and Ping Liu (the Optical Bioimaging Core Facility of WNLO-HUST) for imaging assistance. W.J. was supported by National Natural Science Foundation of China (91854109; 31701170), and the Program for HUST Academic Frontier Youth Team (2018QYTD11). J.X was supported by National Natural Science Foundation of China (81901166).

## Author contributions

Y.D., J.X., and W.J. conceived the project and designed the experiments. Y.D., and J.X. performed the experiments. J Y.D., J.X., and W.J. analyzed and interpreted the data. W.J. prepared the manuscript with inputs and approval from all authors.

## Data and materials availability

All the data and relevant materials, including reagents and primers, that supports the findings of this study are available from the corresponding author upon reasonable request.

---

**Video 1:** 3D reconstructions of TGN-LD interactions were rotating along z-axis in HEK293 cell cultured in OA medium expressing VPS13B^sfGFP (green) and Halo-Rab6 (TGN, blue), stained with BODIPY 558/568 (LD, magenta). A stack of 31 z-planes with 0.2μm thickness in each stack was taken.

**Video 2:** Dynamics of TGN-LD interactions in HEK293 cell expressing VPS13B^sfGFP (green) and Halo-Rab6 (TGN, blue), stained with BODIPY 558/568 (LD, magenta). Time in sec. Scar bar: 2 µm.

**Fig. S1.**
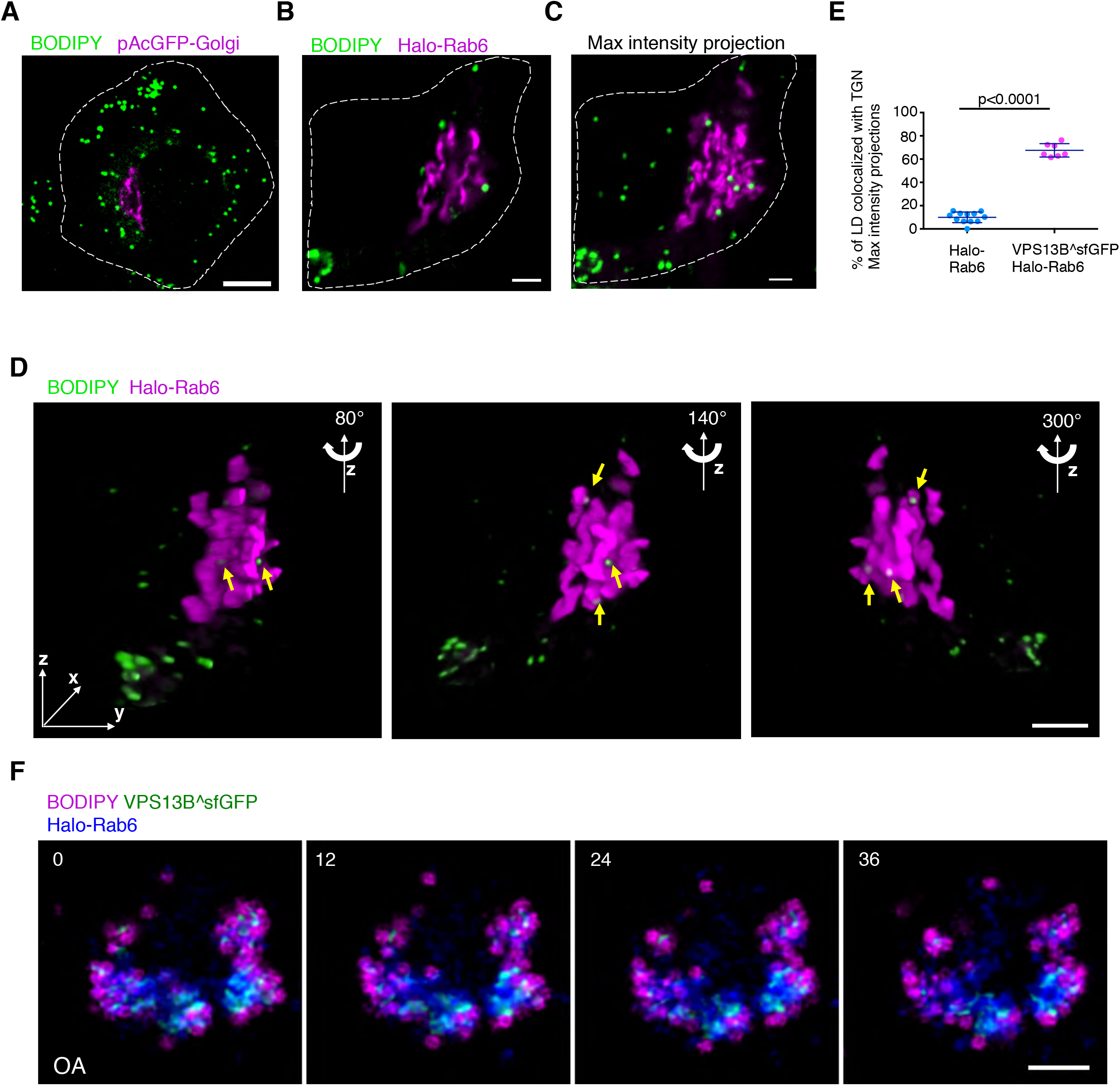
Supplemental data to Fig.1. (A) Representative confocal image of a BODIPY 558/568 (green)-labeled HEK293 cell expressing pAcGFP1-Golgi (magenta). (B) Representative Airyscan image of a BODIPY 558/568 (green)-labeled HEK293 cell expressing Halo-Rab6 (magenta). (C) The maximum intensity projections of 3D reconstructions of the cell in (B) showing TGN-LD interactions. (D) 3D images of the cell in (B) with three different angles (80°, 140°, and 300°) along z-axis were shown. A stack of 26 z-planes with 0.19μm thickness in each stack was taken.. Yellow arrows denoted LDs tightly associating with the TGN. (E) Quantification of TGN-LD interactions in BODIPY558/568-labeled cells either expressing TGN marker pAcGFP-Golgi (n=12) or co-expressing VPS13B^sfGFP and Halo-Rab6 (n=7) based on the maximum intensity projections from 3D constructions. Mean ± SD. Two tailed unpaired student t test. (F) Time-lapse images of a BODIPY558/568 (magenta)-labeled HEK293 cell expressing VPS13B^sfGFP (green) and Halo-Rab6 (blue). Time in sec. Scale bar, 10 µm in (A, B, C & E).

**Fig. S2:**
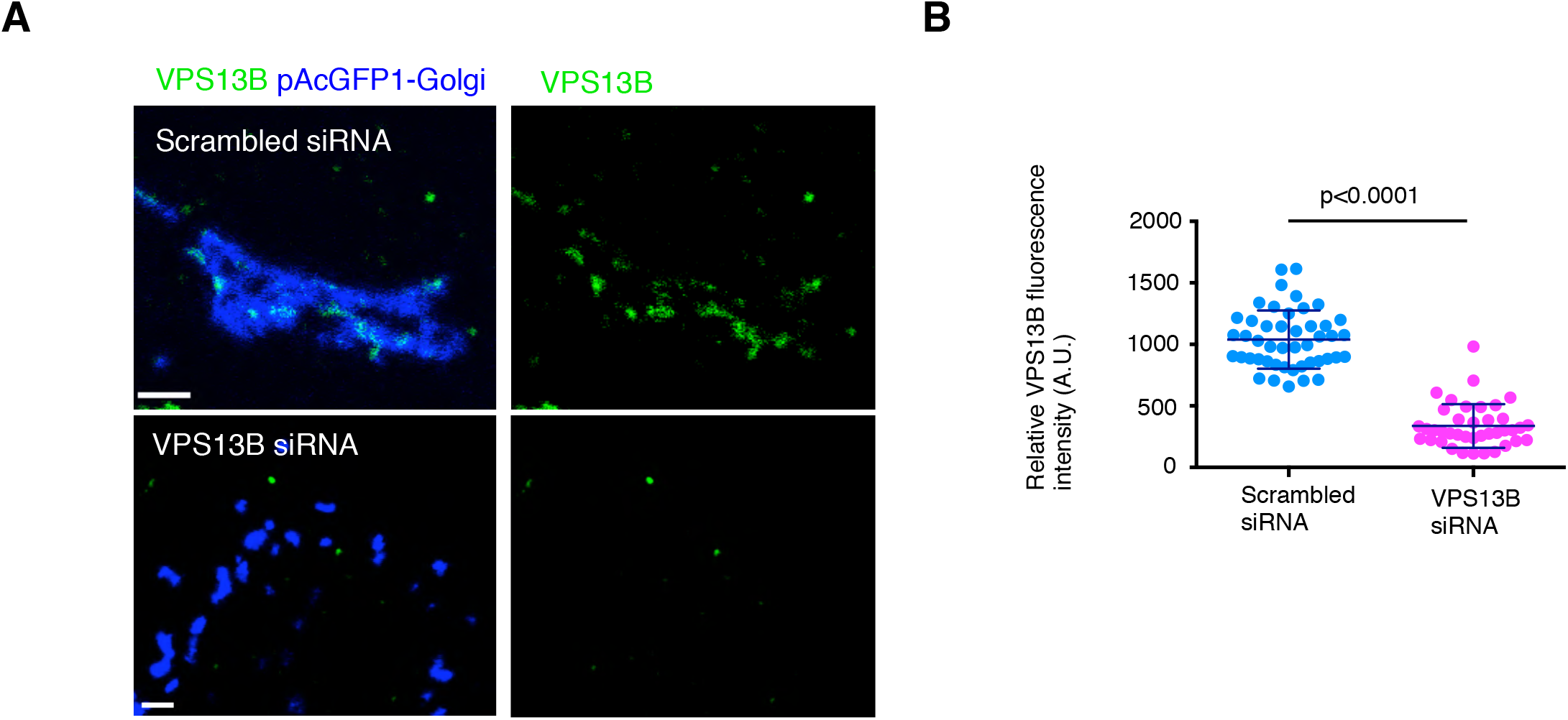
Validation of anti-VPS13B antibody in IF. (A) Confocal images of scrambled (top panel) or VPS13B siRNAs treated (bottom panel) HeLa cells expressing pAcGFP-Golgi (blue) in IF by anti-VPS13B antibody (green). Contrast range of images was set to same level for both cells. (B) Quantification of VPS13B fluorescence intensity in IF of scrambled (n=46) or VPS13B siRNAs (n=39) treated cells. Mean ± SD. Two tailed unpaired student t test.

**Fig. S3:**
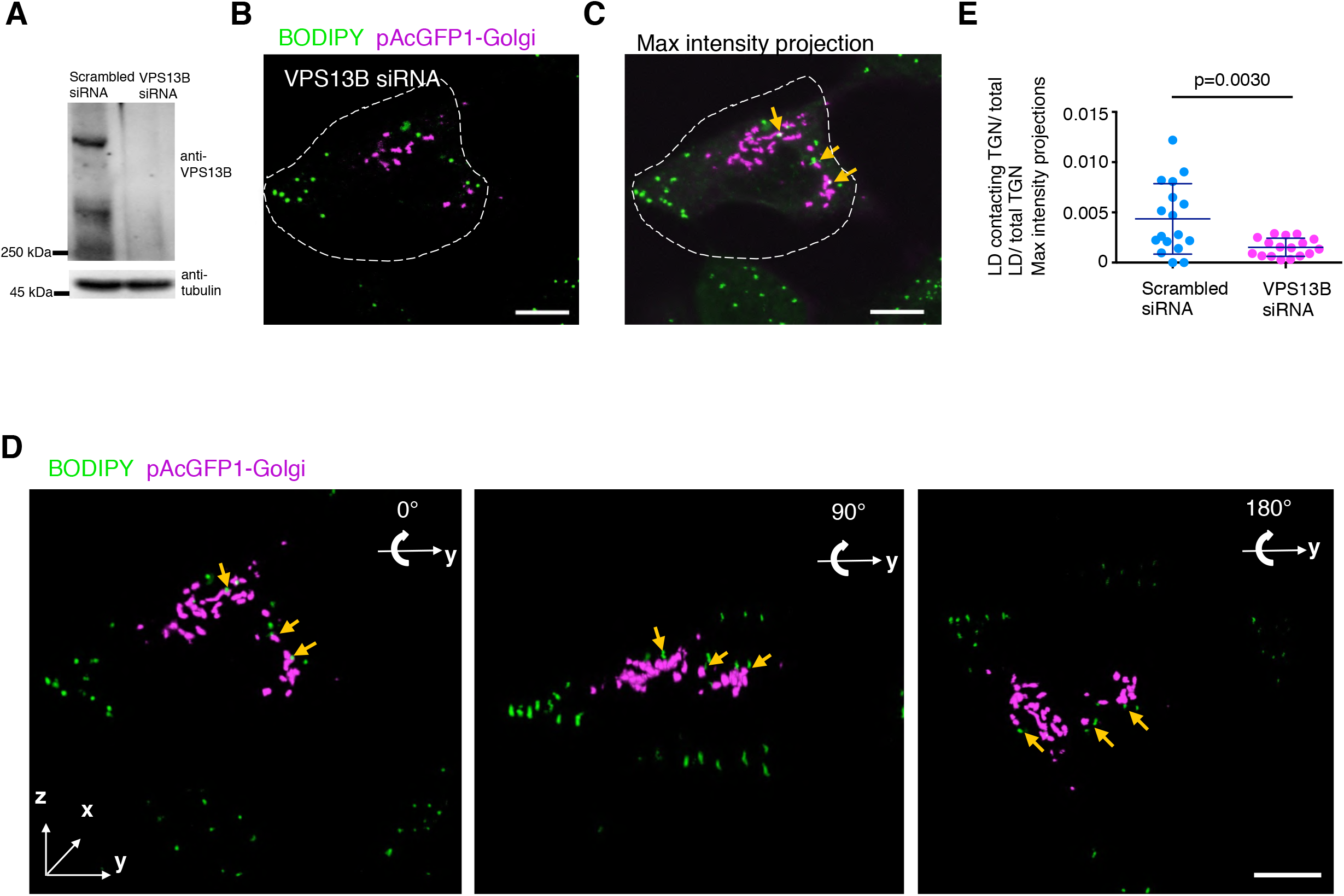
The TGN-LD interactions in VPS13B-depleted cells. (A) Immunoblots demonstrating the efficiency of siRNAs-mediated VPS13B depletion in HeLa cells by using antibody against VPS13B and tubulin. (B) Representative confocal images of a BODIPY 558/568 (green)-labeled HeLa cell expressing pAcGFP1-Golgi (magenta) upon treatments of VPS13B siRNAs. (C) The maximum intensity projections of 3D reconstructions of the VPS13B-depleted cell in (A) showing TGN-LD interactions. (D) 3D images of the VPS13B-depleted cell in (A) with three different angles (0°, 90°, and 180°) rotating along y-axis were shown. A stack of 26 z-planes with 0.19μm thickness in each stack was taken. Yellow arrows denoted LDs adjacent but not physically contacting the TGN. (E) Quantification of TGN-LD interactions in BODIPY558/568-labeled cells expressing pAcGFP-Golgi in response to either scrambled (n=17) or VPS13B siRNAs (n=17) based on the maximum intensity projections from 3D constructions. Mean ± SD. Two tailed unpaired student t test. Scale bar, 10 µm in (B-D).

**Fig. S4:**
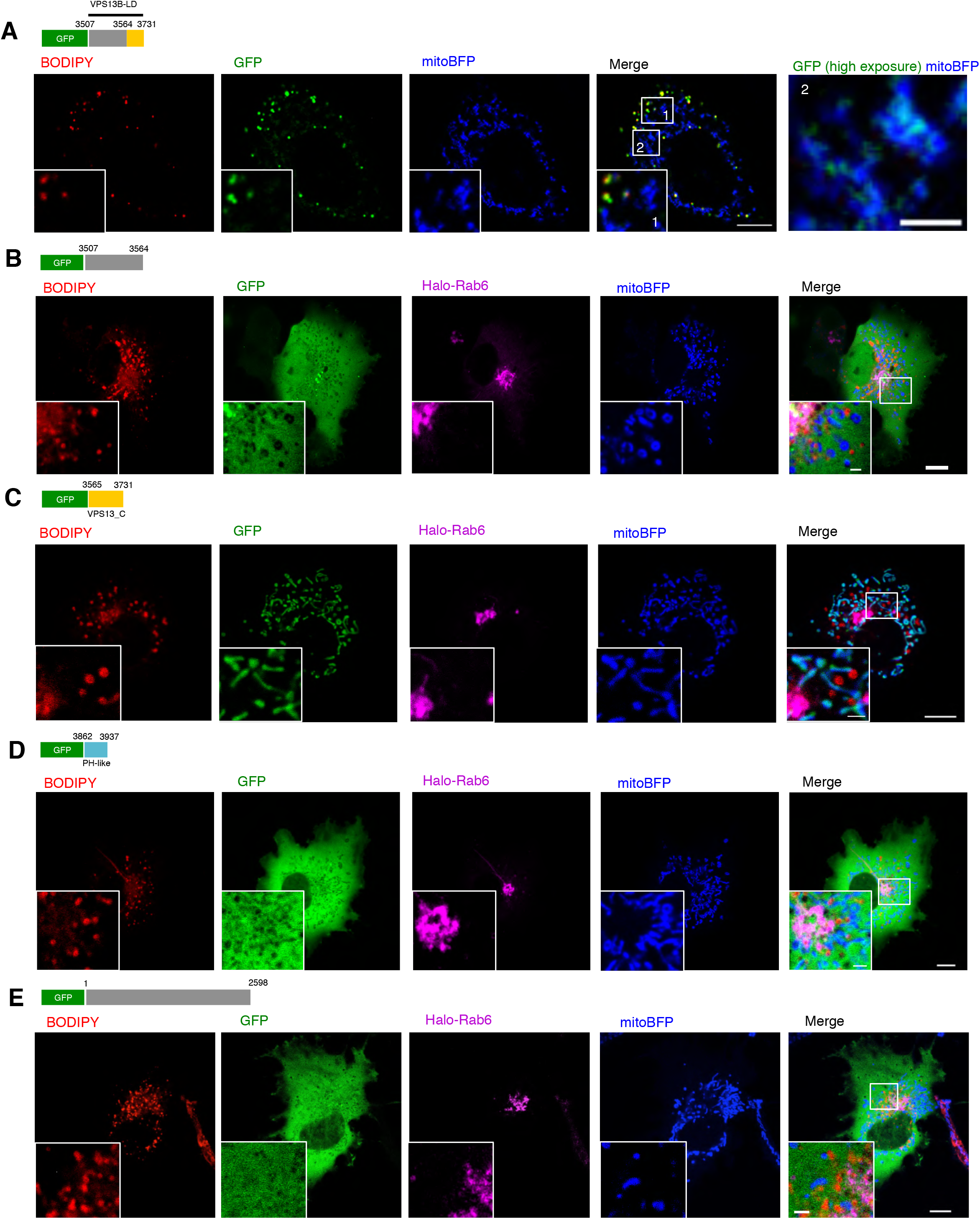
Supplemental data to Fig. 2 and 3. (A) Example of a BODIPY558/568 (red)-labeled HEK293 cell expressing truncated mutant GFP-VPS13B-LD (green), and mitoBFP (blue) with insets on bottom-left (inset-1) and on the right (inset-2, high exposure of GFP fluorescence). (B) Example of a BODIPY558/568 (red)-labeled HEK293 cell expressing a truncation mutant containing residues 3507-3564 (green) and Halo-Rab6 (magenta) with insets on bottom-left. (C) Example of a HEK293 cell as in (B) expressing GFP-VPS13_C (green) with insets on bottom-left showing co-localization between GFP-VPS13_C and mitochondrial matrix marker mitoBFP. (D) As in (B), a HEK293 cell expressing GFP-PH-like (residues 3862-3937) (green). (E) Example of a HEK293 cell as in (B) expressing a truncation mutant containing residues 1-2598 (green). Scale bar, 10 µm in whole cell image and 2 µm in insets in (A-E).

**Fig. S5:**
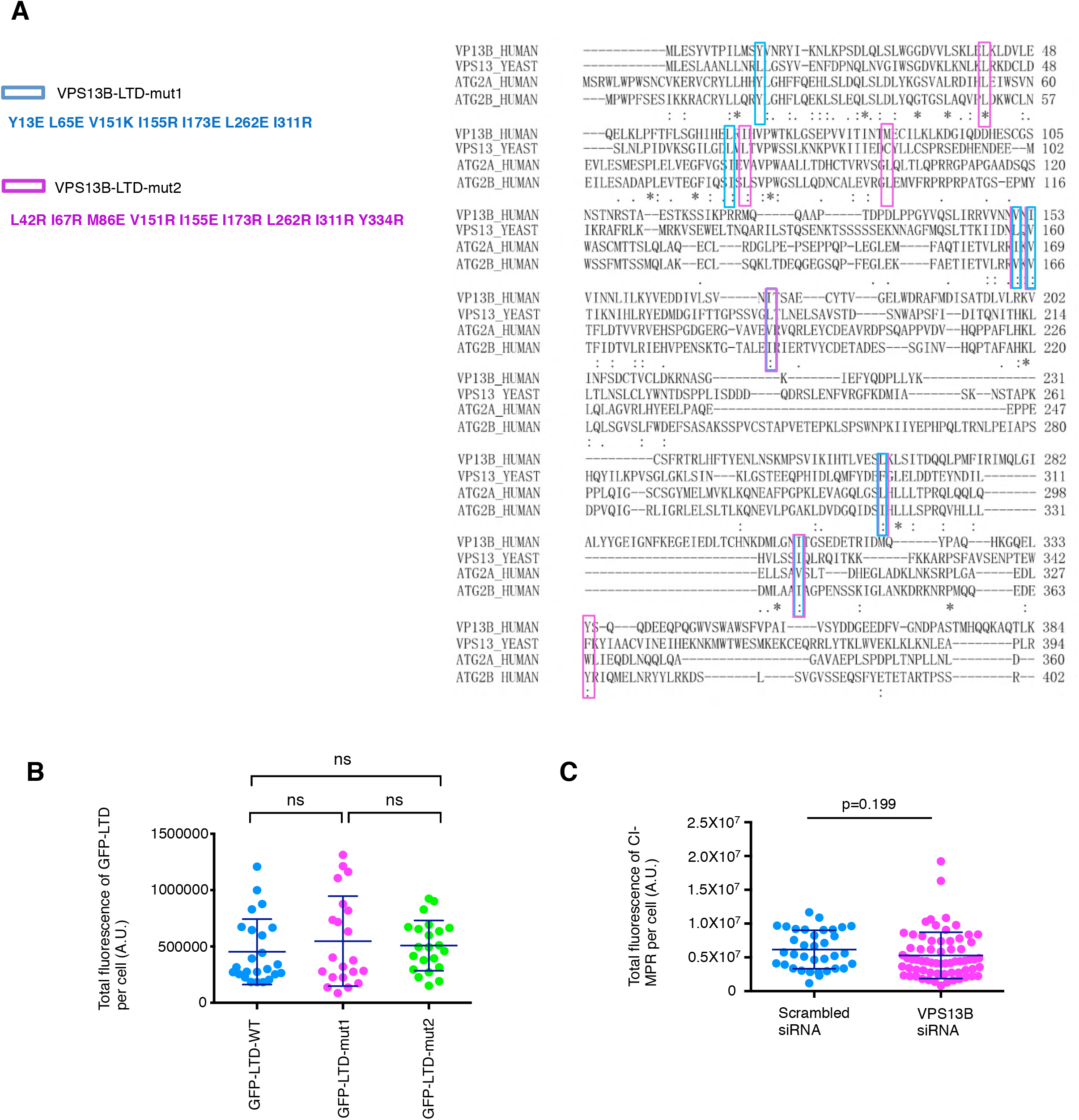
Supplemental data to Fig. 4. (A) Alignment of human VPS13B-LTD (residues 1-384) with human ATG2A-LTD (residues 1-360), human ATG2B-LTD (residues 1-402) and yeast VPS13-LTD (residues 1-360). The conserved hydrophobic residues were mutated to hydrophilic residues, highlighting by either magenta (VPS13B-LTD-mut-1) or blue (VPS13B-LTD-mut-2) box. (B) Quantification of the total fluorescence of GFP-LTD-WT (n=24), GFP-LTD-mut1 (n=21), or GFP-LTD-mut2 (n=22) in HeLa cells in response to VPS13B siRNAs in (Fig. 4F). Not significant (ns) by an ordinary one-way ANOVA followed by Tukey’s multiple comparisons test. Mean±SD. (C) Quantification of the total fluorescence of internalized anti-CI-MPR antibody by immunostaining in scrambled (n=35) or VPS13B siRNAs (n=66) treated HeLa cells. Mean±SD. Two tailed unpaired student t test.

**Fig. S6.**
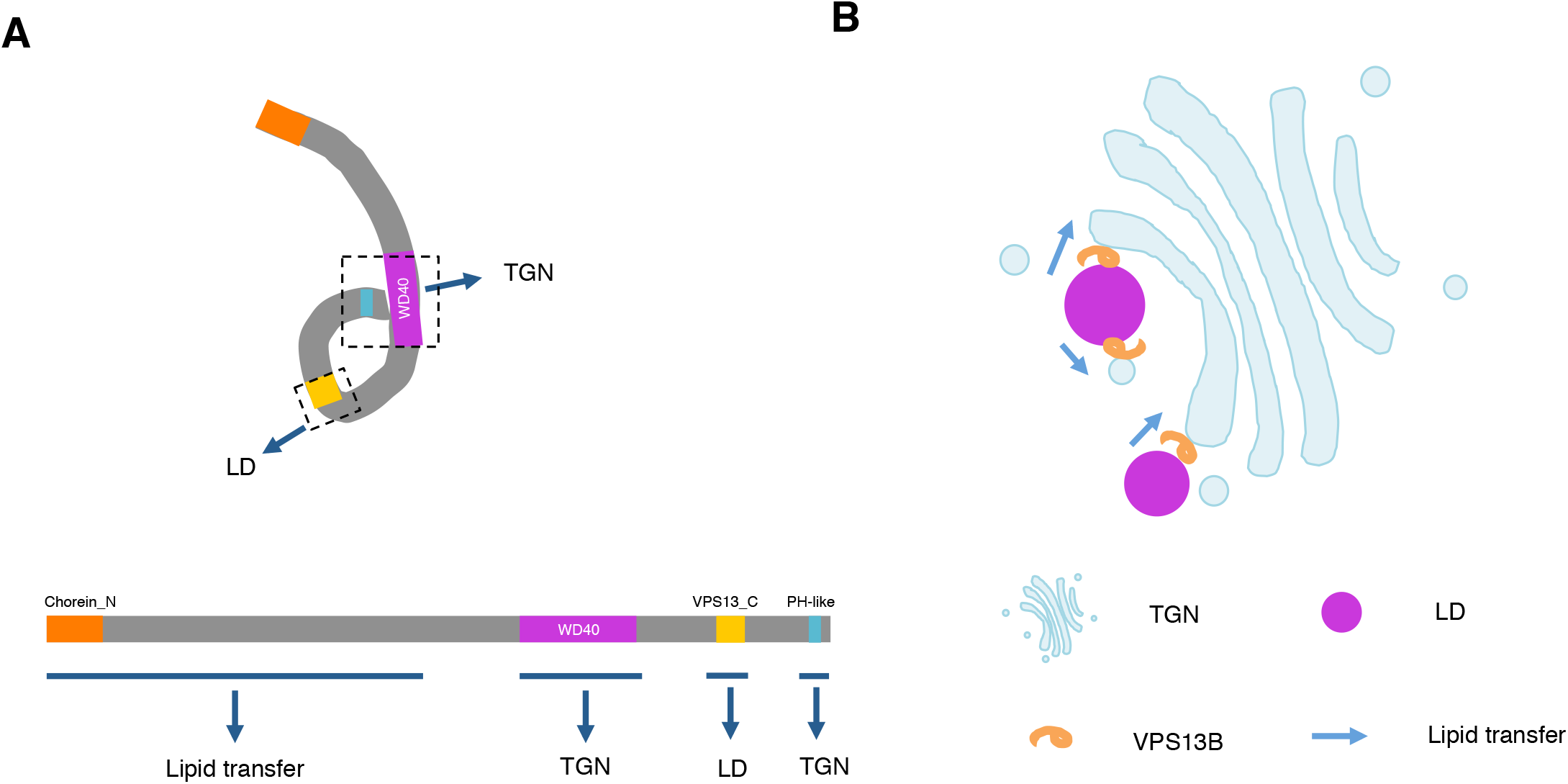
Working model of VPS13B at TGN-LD MCSs. (A) The spatial relation between the domains of VPS13B and its putative protein structure. Not scaled. (B) Models of VPS13B mediated TGN-LD MCSs and lipid transfer activities at such MCSs.

